# A new highly-specific Natural Killer cell-specific gene signature predicting recurrence in colorectal cancer patients

**DOI:** 10.1101/2022.04.29.489868

**Authors:** Carolyn Shembrey, Corina Behrenbruch, Benjamin PT Loveday, Alexander G Heriot, Momeneh Foroutan, Frédéric Hollande

## Abstract

The protective role of Natural Killer (NK) cell tumour immunosurveillance has long been recognised in colorectal cancer (CRC). However, as most patients show limited intra-tumoral NK cell infiltration, improving our ability to identify those with high NK cell activity might aid in dissecting the molecular features which could trigger strong response to NK cell-mediated immune killing. Here, a novel CRC-specific NK cell gene signature capable of inferring NK cell load in primary tissue samples was derived and validated in multiple patient CRC cohorts. The specificity of the signature is substantiated in tumour-infiltrating NK cells from primary CRC tumours at the single cell level, and the expression profile of each constituent gene is explored in NK cells of different maturation states, activation status and anatomical origin. Thus, in contrast with other NK cell gene signatures that have several overlapping genes across different immune cell types, our NK cell signature has been extensively refined to be specific for CRC-infiltrating NK cells and includes genes which identify a broad spectrum of NK cell subtypes. Moreover, it was shown that this novel NK cell signature accurately discriminates murine NK cells, demonstrating the potential applicability of this signature when mining datasets generated from both clinical and animal studies. Differential gene expression analysis revealed tumour-intrinsic features associated with NK cell inclusion versus exclusion in CRC patients, with those tumours with predicted high NK activity showing strong evidence of enhanced chemotactic and cytotoxic transcriptional programs. Furthermore, survival modelling indicated that NK signature expression is associated with improved survival outcomes in two large cohorts of primary CRC patients. Thus, scoring CRC samples with this refined NK cell signature might aid in identifying patients with high NK cell activity who could be prime candidates for NK cell directed immunotherapies.

## Introduction

Despite improvements in surgical oncology and precision medicine, 5-year survival rates for patients with late-stage colorectal cancer (CRC) remain extremely poor and innovative treatment strategies are needed. Although T-cell directed immunotherapies are strikingly effective in many solid cancer types, durable responses are limited to approximately 5% of all CRC patients (1). Downregulation of major histocompatibility complex class I (MHC-I), whose expression is a pre-requisite for T-cell-mediated immune killing, has been reported as a major cause of secondary resistance to checkpoint blockade therapy (2,3). Thus, there is an urgent need to explore alternate immunotherapeutic strategies which harness other antigen-independent cytotoxic cell types such as NK cells.

In recent years, natural killer (NK) cells have emerged as a promising candidate for immunotherapeutic targeting. Adoptive transfer of primary NK cells suppresses T-cell mediated graft-versus-host disease and exacerbates graft-versus-tumour responses (4–6). Similarly, a myriad of alloreactive NK cell lines - primarily NK92, but also NKL, KHYG01 and YTS – have demonstrated safety and efficacy in HLA-mismatched recipients in early-stage clinical trials (reviewed in (7). This suggests that NK cell therapeutics may be harnessed in an “off-the-shelf” manner in the future, circumventing the difficulty associated with the *ex vivo* expansion of antigen-specific T-cell clones. Likewise, chimeric antigen receptor (CAR) NK cells targeting CD19 (8,9)and the epidermal growth factor receptor (EGFR; 10,11) have shown promising results in xenograft models, and monoclonal antibodies which neutralise NK cell inhibitory receptors such as the KIR family (12,13) and NKG2A (14) have entered early-stage clinical trials.

NK cell activity is inversely correlated with cancer incidence (15) and there is a wealth of evidence supporting the role of NK cells in controlling both spontaneous and experimental metastasis (16–20). In CRC, NK cell infiltration has been identified as a positive prognostic marker in both primary (21,22) and metastatic (22,23) disease. NK cells differ from conventional lymphocytes in that they function in an MHC-I unrestricted manner in accordance with the “missing-self” hypothesis (24,25). In this manner, NK cell-directed immunotherapies may overcome the restricted benefit of antigen-specific T-cell responses in tumours with high mutational diversity. Moreover, harnessing NK cell cytotoxicity is a promising opportunity in the treatment of immunologically “cold” tumours such as CRC that undergo loss of MHC-I expression. Indeed, recent studies have reported that 40% of patient-derived CRC organoids exhibit MHC-I loss which could not be rescued by IFN stimulation (26), concordant with clinical reports of MHC-I loss in approximately 60% of MSI CRCs (27). Yet, clinical enthusiasm has been tempered due to the scant NK load reported in most CRC tumors, despite high levels of chemokines and cytokines (28). Thus, novel means of determining which patients show high NK cell infiltration and activity and might therefore benefit from NK cell directed immunotherapies is needed.

Transcriptomic signatures are sets of genes whose coordinated expression has a verified association with specific biological parameters – such as cell type and phenotypic state – or clinical measures, including disease subtype, survival outcome or therapeutic response. Deconvoluting the relative abundance of immune cell subsets using transcriptomic signatures offers many benefits over the comparatively low-throughput methods of immunophenotyping which are currently available. In CRC alone, a multitude of prognostic signatures have been reported (29–32) alongside several signatures which predict response to 5-FU based chemotherapies in Stage II-III disease (33–35). Additionally, RNAseq data from a cohort of 40 CRC patients with metastatic or relapsed disease was used to derive a 27-gene signature able to discriminate responders and non-responders to the FOLFOX6 chemotherapy regimen with accuracy of 92.5%, demonstrating the powerful role which signature analysis may play in personalising patient care (36).

Indeed, immunophenotyping of tumour-infiltrating immune cells via multiparametric flow cytometry relies on a limited number of phenotypic markers whose expression may be perturbed by cells in differing maturation or activation states. Notably, NKp46 (encoded by NCR1), an archetypical marker of NK cells, shows variable expression across NK cell subsets (37) and can be significantly downregulated in NK cells from cancer patients (38). Similarly, whilst immunohistochemical approaches provide an unrivalled insight into the spatial topography of immune cell infiltration at the protein level, the limited capacity for co-detection of multiple markers on individual cells limits our ability to accurately identify specific immune cell subsets or activation states in the tumour microenvironment (TME). In their simultaneous assessment of multiple markers, transcriptome-wide approaches circumvent the limitations of single-marker or low-throughput phenotypic assessments. This may also allow for previously unidentified markers or associations to be identified which may provide important insights into immune cell biology.

However, a major consideration when employing such computational approaches is the suitability of the reference profiles, as the gene signatures derived from sorted cell types in healthy individuals may not accurately reflect those of the potentially dysregulated cells in a diseased individual. This is particularly pertinent in the case of NK cells, as activated NK cells have unique transcriptional profiles as compared with resting NKs from healthy volunteers (39). Additionally, as signatures often rely on highly expressed genes and thus often incorporate genes that are not that specific of a given cell type, the accuracy of cell type detemination may be compromised. Although prognostic NK cell gene signatures exist for highly immunogenic tumour types such as melanoma (40) and renal cell carcinoma (41), it is currently unclear how appropriate these are for use in tumours such as CRC which traditionally show a poor immune cell infiltration. Other works which have looked at the transcriptomic profiles of lymphocytes in CRC have focussed on defining residency versus exhaustion states (42), where the genesets used to define CD4+ and CD8+ T-cells had several genes overlapping with the NK cell geneset.

Here, we present a novel NK cell transcriptomic signature which can be used to infer NK cell abundance from bulk RNA sequencing data of primary CRC samples. Candidate NK cell-related genes were pooled from previously published works and in-house differential expression (DE) analyses and sequentially filtered to ensure their fidelity as NK cell markers with minimal off-target expression in tumour, stromal and other immune cell types. Single-cell RNAseq (scRNAseq) data from primary CRC samples were then used to validate each signature gene in tumour-infiltrating NK cells. We then show that high NK cell score is associated with upregulation of cytolytic and chemotactic transcriptional processes, and survival analysis revealed that patients with higher evidence of NK cell activity demonstrate significantly longer recurrence and disease-free intervals. Collectively, the NK cell signature allows for the identification of CRC patients with high NK cell activity, which may aid in defining the molecular characteristics associated with strong response to NK cell targeting immunotherapies.

## Methods

### Preparation of publicly available RNAseq data

Datasets used in the present study are listed in Table S4. Raw counts files from publicly available datasets were downloaded rom Gene Expression Omnibus (GEO) using the NCBI portal (http://www.ncbi.nlm.nih.gov/geo/). CCLE data was downloaded as a PharmacoSet (PSet) through the PharmacoGx R/Bioconductor package (version 1.6.1). For *In-house data*, 3’ RNA-seq reads were aligned using HISAT2 against human genome GRCh37 (release 75) and the featureCounts tool from the RSubread package was used to quantify the number of reads for each gene per sample. The filterByExpr function from the edgeR package (v 3.28.1) was used to filter lowly expressed genes and calculated count- or transcript-per-million (CPM/TPM) values. The MyGeneset application of the ImmGen online databrower (http://www.immgen.org/Databrowser19/DatabrowserPage.html) was used for analysis of GSE15907.

### Differential expression analysis, gene set testing and survival modelling

The singscore (v1.6.0) R package was used for Single-sample gene set enrichment analysis against various molecular signatures. Depending on the direction of gene sets, we used different settings of simpleScore function as specified in the documentation. NK-high and NK-low groupings were defined as the top and bottom 10% of samples, respectively, when ranked by NK scores. DE analysis between NK-high and NK-low samples was performed using the voom-limma (v 3.42.2) pipeline (Law *et al*., 2014). After running eBayes, we considered genes with absolute log_2_FC > 1 (for ovexpression) or log_2_FC < −1 (for repression) and adjusted p-value < 0.05 as DEGs. The goana function from the limma package was used to perform gene ontology (GO) analysis and camera gene set testing was performed using MSigDB signatures retrieved from the WEHI bioinformatics portal (http://bioinf.wehi.edu.au/software/MSigDB/). Survival modelling was performed using the survminer (v0.4.8) and survival (v3.2-7) packages using the clinical annotation files provided from the sources listed in Table 2.14.

### Data wrangling and visualization

All computational analyses were performed using R (version 3.6.1). For data wrangling and visualization, base R functions were using alongside several core packages from the tidyverse (v 1.3.0) R package. tidyr (v 1.1.2) and dplyr (v 1.0.2) were used for reading and manipulating the data, as well as ggplot2 (v 3.2.1) and cowplot (v 1.0.0) for data visualization. Heatmaps were generated using the complexHeatmap (v 2.2.0) or pheatmap (v 1.0.12) R packages.

### Statistical Analysis

All statistical analyses were performed using R. Data is expressed as mean ± standard deviation (SD) unless otherwise indicated. The minimum threshold for rejecting the null hypothesis was p<0.05. For results where statistics are shown, significance is denoted as: * = p<0.05; ** = p< 0.01; *** = p<0.001.

### Code availability

The code used throughout this study will be made available on Github upon publication

## Results

### Collation of candidate NK cell signature genes

To identify a CRC-specific gene expression signature associated with NK cell abundance, 609 unique putative NK cell-associated genes were collated from eight partially overlapping sources (Figure 1). Firstly, four previously curated NK cell signatures were compiled. “CIBERSORT Active” (n_genes_ = 56) and “CIBERSORT Resting” (n_genes_ = 56) refer to the gene sets corresponding to activated and resting NK cells, respectively, as reported in the LM22 signature matrix used for the CIBERSORTx algorithm (43). The “Cursons Extended NK cell Geneset” gene set (n_genes_ = 112 genes) was derived from the supplementary data table of a melanoma-specific NK cell signature previously reported by Cursons and colleagues (40). The “Wang NK cell marker” gene set (n_genes_ = 13; Wang *et al*., 2017a) is composed of markers used to guide the immunophenotyping of different cell subsets in bulk RNAseq data from CRC cell lines and primary samples, and the “receptors” gene set (n_genes_ = 43) was compiled by mining the literature for various receptor subsets (eg. activating, inhibitory, chemokine or cytokine receptors) with documented expression on NK cells.

**Figure 1.**
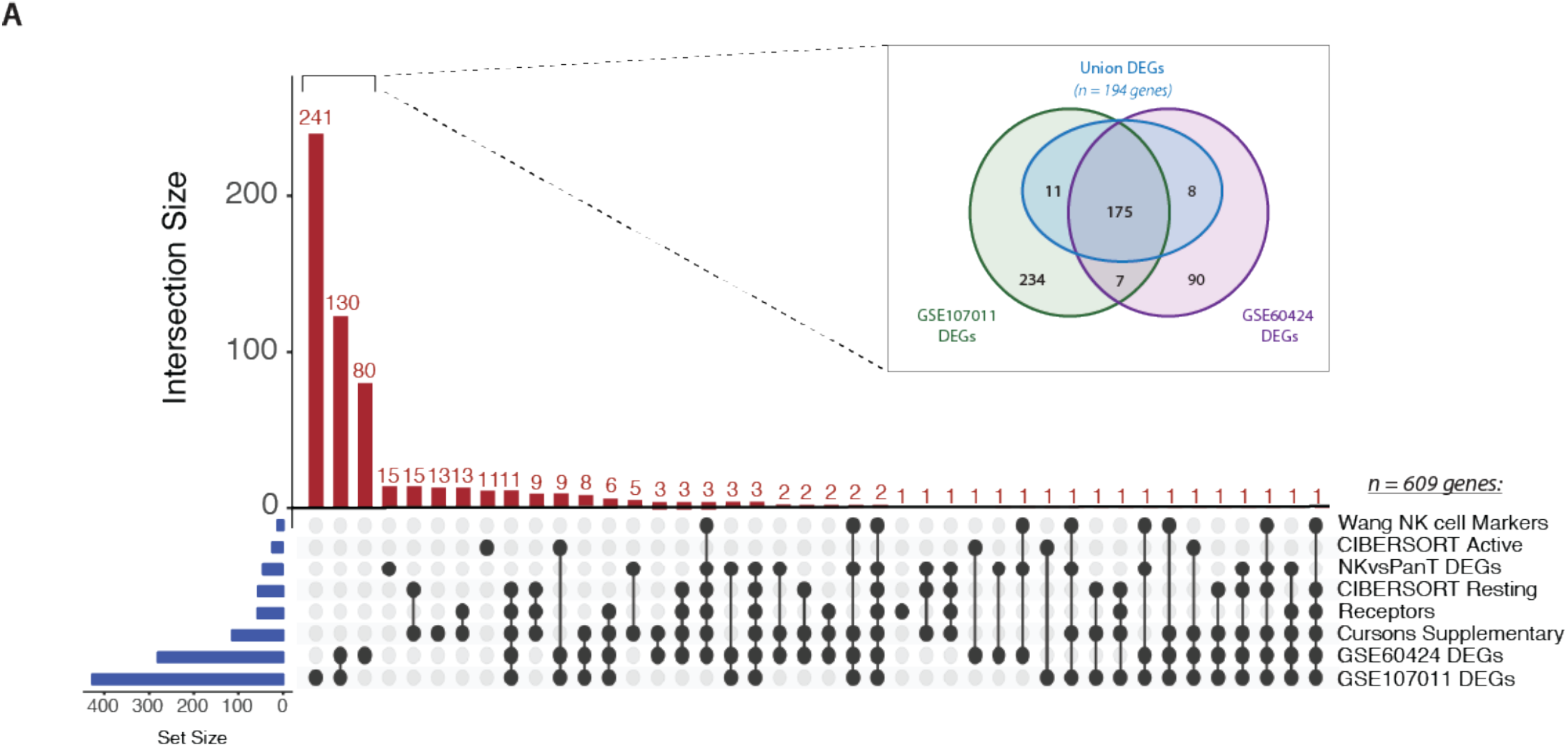
Collation of putative NK cell signature genes. UpSet plot of gene intersects from eight partially overlapping sources (listed at right). Gene set (blue bars) and intersection (redbars) sizes are indicated. Inset: Venn diagram of DEGs identified from pairwise comparisons of NK cells versus other immune cell types in GSE60424 and GSE107011. Only the “UnionDEGs” (ngenes= 194) were retained as candidate signature genes

Subsequently, three novel gene sets were derived from the results of differential expression (DE) analysis, where putative genes enriched in NK cells were identified by pairwise comparison of NK cells and at least one other immune cell type. DE analysis of the GSE60424 bulk RNAseq dataset (44), composed of six immune FACS-isolated cell types (NK cells, B-cells, CD4+ T-cells, CD8+ T-cells, monocytes and neutrophils), identified 280 DEGs (“GSE60424 DEGs”). Similarly, DE analysis of GSE107011 (45), an RNA-seq dataset composed of 29 FACS-isolated immune cell types, yielded 427 DEGs (“GSE107011 DEGs”). Finally, to enhance our resolution when discriminating between NK and T-cells, a subsequent “pan-T DEGs” (n_genes_ = 23; from GSE107011) was constructed from the DE genes when only NK versus T-cell comparisons were considered; here, the selected genes were those upregulated in NK cells relative to all T-cell subsets (eg. upregulated in NKs relative to CD8+ *and* CD4+ *and* MAIT etc.), irrespective of their expression level in other innate immune cell types.

Following compilation, 609 unique candidate NK genes were identified. To examine the representation of traditional NK cell markers versus potentially novel NK cell-related genes, the interconnectedness of this gene set was compared across the eight sources (Figure 1). Most genes identified (451/608; 74%) were uniquely found from our DE analyses. Strikingly, there were zero genes which were conserved across all eight sources and only two genes, KLRD1 and KIR3DL2, were conserved across 7/8 sources.

To further refine the comparatively large GSE60424 and GSE107011 DEG sets, a “Union DEGs” gene set was created (Figure 1; inset). For inclusion in this merged gene set, a given candidate gene needed to be differentially upregulated in NK cells relative to all other immune cell types in one or both data sets (eg. upregulated in NK cells versus T-cells *and* B-cells *and* neutrophils) differing from the previous DEG analyses where the gene in question need only be upregulated in NK cells on a pairwise basis (eg. upregulated in NK cells versus T-cells, but not necessarily B-cells nor neutrophils). Of the 525 DEGs identified across GSE60424 and GSE107011, 194 genes fit this criterion; 175 were present in both data sets, with an additional 11 genes being specific to GSE107011 and 8 genes specific to GSE60424. Whilst a further 7 genes were commonly identified as DEGs in both data sets, these genes were not upregulated in NK cells relative to all other immune cell types and therefore excluded from the “Union DEGs” set. After reintegration of these “Union DEGs” with the genes derived from other sources, 295 candidate genes remained.

Although DE analysis identifies genes which are preferentially expressed by NK cells as compared with other cell types, the expression of individual DEGs by NK cells may still be very low. This is problematic when aiming to identify NK cell signals from tumour sequencing data, particularly given that NK cells represent a small proportion of the total cell number. Thus, we further refined out geneset by retaining only those candidates with higher median in NK cells relative to other immune cell subsets in GSE107011 (Figure 2A, Figure S2A; purple boxes) and GSE60424 (Figure S2B; purple boxes). Sets of passing genes for each dataset were derived by taking the intersect of the specific genes (n_genes_ = 49) from GSE107011 (Figure 2A) and GSE60424 (Figure S2B). Of these, 10 genes (ADGRB2, B4GALT6, LDB2, LIM2, LINC01451, LRRC43, PCDH1, PRSS57, RAMP1 and RNF165) were identified in both data sets.

**Figure 2.**
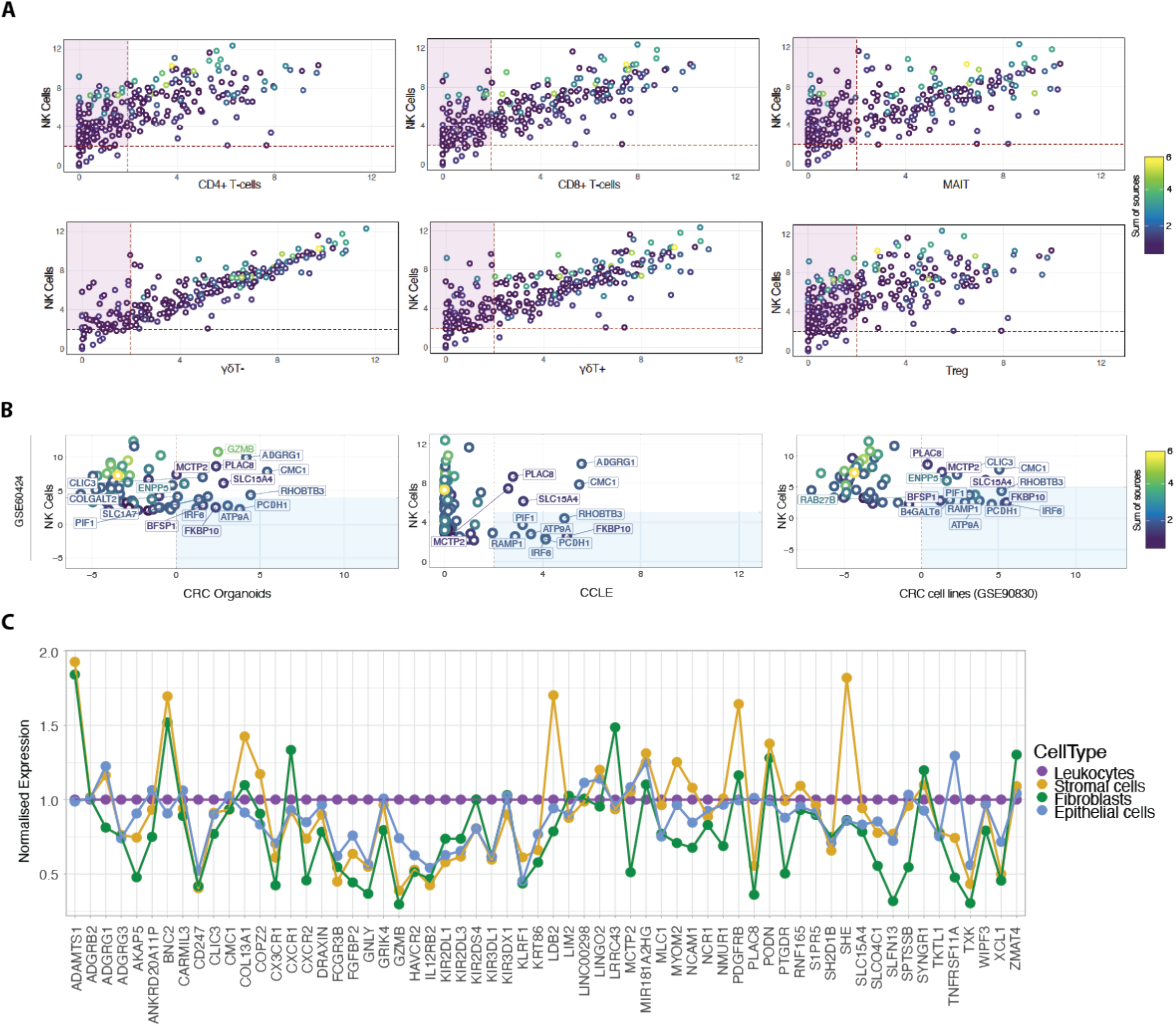
NK cell specificity filtration against immune and CRC-supportive cell types. (A) Biplots depicting median expression for each candidate gene (rings; coloured by sum of sources) in NK cells from GSE107011 (logTPM) versus other immune cell types. The intersect of passing genes for each pairwise comparison (purple boxes) were retained as candidate genes. (B) Biplots depicting median expression for each candidate gene (rings; coloured by sum of sources) in NK cells from GSE107011 (logTPM) versus CRC organoids(left column; logRPKM), CCLE CRC cell lines (centre column; logRPKM) and GSE90830(CRC cell lines; right column; logRPKM). The union of failing genes for each pairwise comparison (blue boxes; ngenes= 12) were flagged for removal from the candidate geneset. (C) Line plot of candidate gene expression in the stromal cell (CD31+; gold), fibroblast (FAP+; green) and epithelial cell (EpCAM+; light blue) compartments of six primary CRC samples (GSE39396), normalised to leukocyte (CD45+; purple) expression. Genes with normalised expression >1 were flagged for removal. CCLE: Cancer Cell Line Encyclopedia; CoT: CRC primary tumour; LT: CRC liver metastasis

Interestingly, this analysis highlighted that many of the “classical” NK cell markers that are abundant in multiple sources (Figure 2A; yellow/green rings) are also expressed at very high levels in “unconventional” T-cell subsets such as MAIT, γδ+ and γδ-T-cells. As illustrated by GSE60424 (Figure S2B), where T-cells are exclusively grouped as CD8+ or CD4+, many FACS-based studies do not include these relatively niche T-cell subsets, possibly leading to the identification of putative NK cell-specific genes which are in fact highly expressed by unconventional T-cell subsets.

The two sorted NK cell-containing data sets interrogated thus far (GSE60424 and GSE107011) were derived from the peripheral blood of healthy individuals; this approach has limitations, as the gene expression profiles of such cells may not necessarily reflect those of tissue-infiltrating NK cells nor NK cells in the context of cancer. To address this concern, we performed an independent DE analysis (see methods for details) on an NK-cell containing scRNAseq data set (GSE146771, (46), composed of SMART-Seq2 and 10X subseries) generated from a cohort of CRC patients (n_samples_ = 20; from 18 unique patients) with Stage IIIII disease of varying pathological grade, MSI status, and extent of nodal involvement. This analysis identified several additional NK cell marker genes, and also allowed us to crossvalidate the 49 genes derived from bulk RNA-seq analysis (mentioned above) at the single cell level. Following these analysis steps, 82 candidate genes were prioritised.

### Signature gene specificity filtration against tumour and stromal cells

As the NK cell signature is designed to resolve the NK cell fraction from bulk RNAseq data of tumour tissue, it is imperative that the genes in the signature should not be expressed by the tumour cells themselves. To determine whether any of the 82 identified genes were expressed by tumour cells, additional specificity thresholding was performed against three sources of CRC cells (Figure 2B). As CRC tumours themselves were expected to contain immune cell fractions, thereby confounding any interpretations, *in vitro* cellular models of CRC were used in this analysis. RNAseq data from two partially-overlapping repositories of CRC cell lines were included, The Cancer Cell Line Encyclopedia (47) n_samples_ = 57) and GSE90830 (n_samples_ = 44 (39)), alongside RNAseq data from an array of patient-derived organoids (PDOs; (n_samples_ = 21, (40)) generated in-house from primary CRC and colorectal liver metastasis (CRLM) tissues. Interrogation of our candidate gene list in these CRC datasets revealed that 12 genes had relatively higher expression in CRC cells compared with NK cells (Figure 2B; blue boxes), and they were therefore excluded from further analyses.

Despite the immune- and tumour-specific filtration steps previously performed, the possibility of contaminant expression of our candidate genes by other non-NK cell types such as including fibroblasts, stromal cells and non-transformed epithelial cells has not been accounted for. To address this, the relative expression of our candidate genes was interrogated in the leukocyte (CD45+), stromal cell (CD31+) and epithelial cell (EpCAM+) fractions isolated from the tumours of 6 patients with CRC (GSE39397; (50)as well as against cultured, normal colon mucosa-derived fibroblasts (CCD-Co-18). Failing genes (n_genes_ = 14) were defined as those with significantly higher expression relative to leukocytes in a non-leukocyte subset (Figure S3; red headers, summarised in Figure 2C).

### Interrogating expression of signature genes in tumour-infiltrating NK cells

Having validated the specificity of our candidate genes at the immune, tumour and stromal cell level in bulk RNAseq, we returned to GSE146771 to confirm this specificity in scRNAseq data. By interrogating the gene expression distribution across Uniform Manifold Approximation and Projection (UMAP) plots (Figure 3), the specificity of each gene for particular NK cell subsets was elucidated. For example, in the SMART-Seq dataset three NK cell subgroups could be discerned based upon marker expression, providing greater cellular resolution: CD16^+^, a marker expressed on virtually all CD56^dim^ NK cells; GZMK^+^, a cytotoxicity marker frequently associated with the CD56^bright^ subpopulation; and CD103^+^, a marker of tissue residency. Certain “NK cell specific” genes such as SH2D1B and KIR2DL3 showed equivalent expression in each of the three NK subsets, whereas other genes were subtype selective. For example, LINGO2, PRSS57 and RNF165 were preferentially expressed by the CD16+ subgroup, whereas WIPF3 expression was high across both the CD103+ and GZMK+ subgroups but sparse in the CD16+ population.

**Figure 3.**
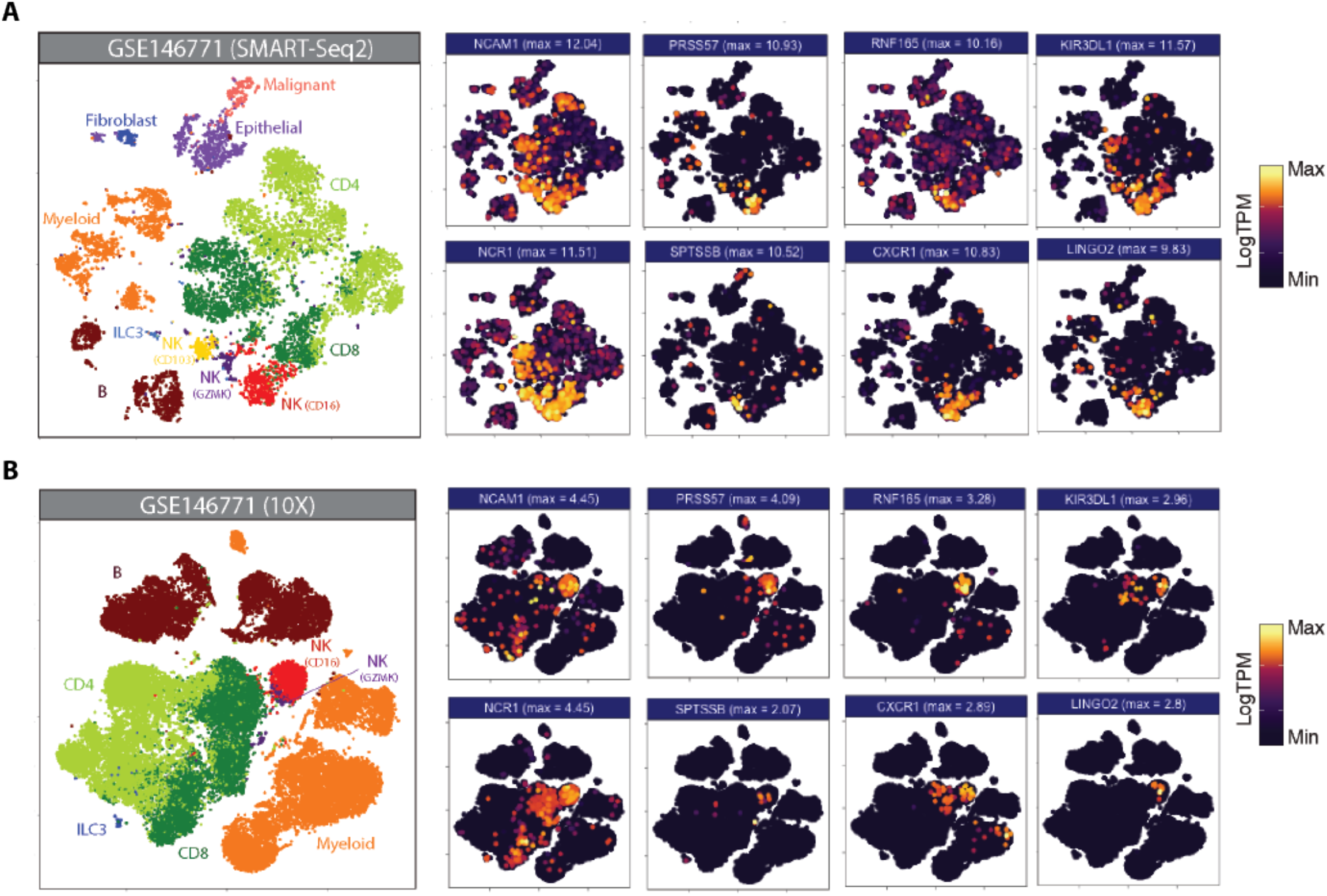
Candidate gene expression in CRC-infiltrating immune cells. UMAP plots of dissociated primary CRC samples from (A)GSE146771 (SMART-Seq2 scRNAseq; n samples = 10) and(B)GSE146771 (10X scRNAseq; n samples= 10) coloured by cell type (at left) and candidate gene expression. Maximum expression (LogTPM) is indicated in parentheses above each plot.

This approach also allowed us to identify genes which, although differentially upregulated in NK cells, showed moderate basal expression across multiple cell types (Figures S4, S5; “High DEGs” pale blue headers) as well as genes whose expression was no longer specific to NK cells once evaluated at the single-cell level (Figures S4, S5; “Non-specific”; red headers). For example, many candidate genes - including PLAC8, SLC15A4, SLFN13 and ST8SIA6 – have high expression in NK cells although they are promiscuously expressed by multiple cell types, with particularly high expression in the myeloid subset, warranting their exclusion from the final gene list.

### Validating the subtype specificity of the curated NK cell signature

Having refined the NK cell signature using CRC-infiltrating NK cells, we next interrogated whether the signature exhibited any biases towards NK cells from particular subsets or sources. GSE133383 (37) contains transcriptomic data for both the immature CD56^bright^ and mature CD56^dim^ subsets of NK cells isolated from the blood, lymphoid organs (spleen, lymph nodes and bone marrow) and lungs of four healthy donors. As is the case with most transcriptional analyses of NK cells, when interrogated in GSE133383 our signature genes cluster according to NK cell maturation state rather than tissue source (Figure 4A). Whilst a substantial number of genes which appear to have relatively uniform expression across both the CD56 bright and dim subsets (Figure 3.7A and Figure S4), multiple subtype-enriched markers have also been retained. Enriched markers for the populous CD56^dim^ class include the KIR family members, GZMB (37,51,52) and the chemokine receptors CXCR1 and CXCR2 (53). Two recently identified markers of terminally mature NK cells, HAVCR2 and CXC3CR1 (54) show preferential expression in the CD56^dim^ subset as expected. Analogously, enriched markers for the CD56^bright^ subset include CD56 (NCAM1) itself, XCL1 (54,55) and KLRC1 (56), whose inclusion indicate that the signature has sufficient resolution to detect this relatively minor population. Collectively, this demonstrates that our signature captures both general and subset-enriched NK cell markers, suggesting that it is representative of all types of NK cell.

**Figure 4.**
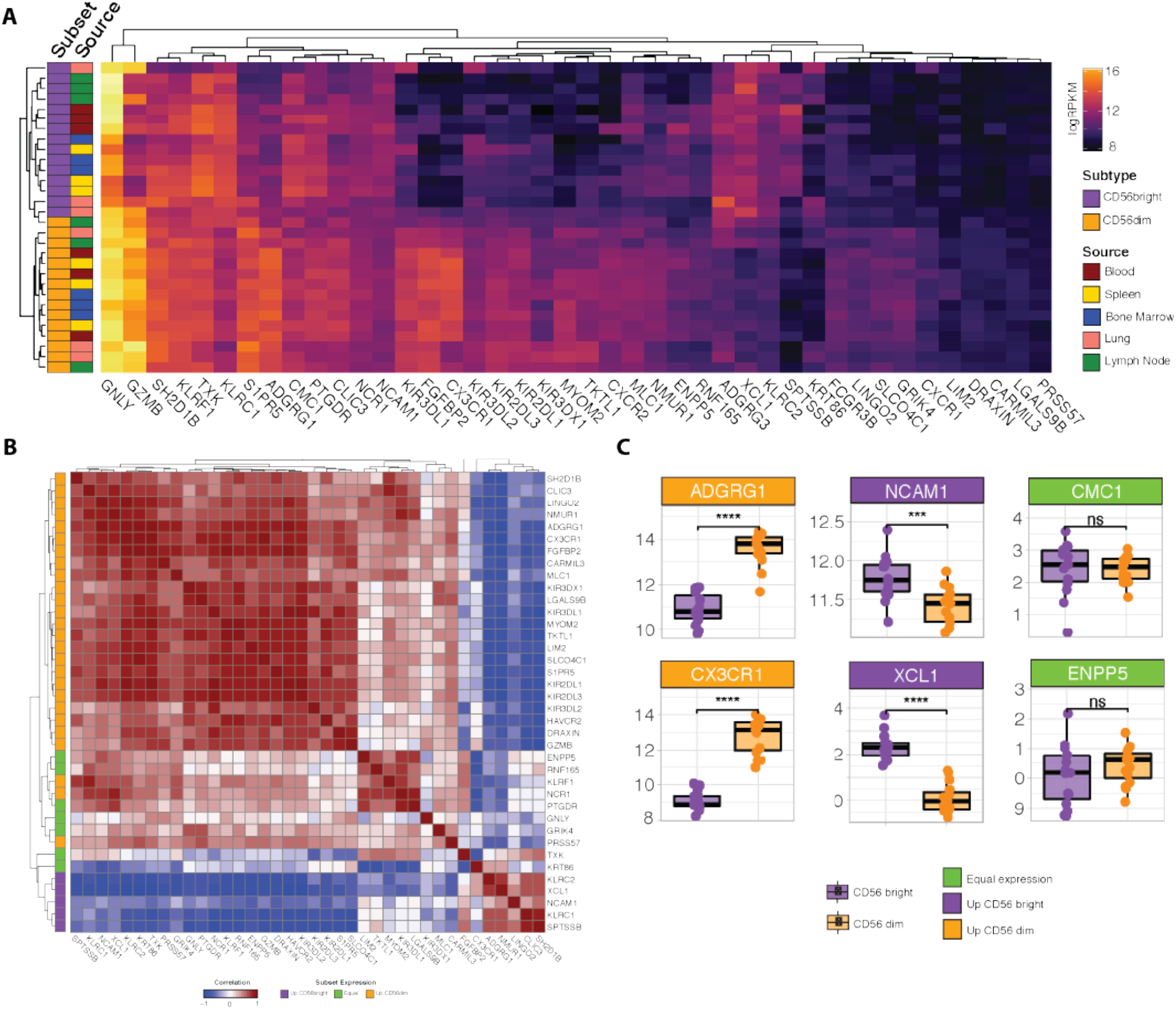
Profiling refined gene set expression in NK cell subsets. **(A)** Heatmap of refined gene set expression in CD56^bright^ (orange) and CD56^dim^ (purple) NK cells isolated from the blood (red), spleen (yellow), bone marrow (blue), lung (pink) and lymph node (green) of four healthy donors (GSE133383). **(B)** Correlation matrix of refined gene set. Genes are annotated (at left) according to whether they show preferential expression in the CD56^dim^ (orange), CD56^bright^ (purple) or neither (green) NK cell population **(C)** Boxplot expression of selected genes in CD56^dim^ (orange boxplots) vs CD56^bright^ (purple boxplots) NK cell populations. Headers are coloured based on whether the gene is preferentially expressed in CD56^dim^ (orange headers) vs CD56^bright^ (purple headers) NK cells, or shows equivalent expression across both subsets (green headers). Significance was assessed with Student’s T-test; *: p-value < 0.05; **: p-value < 0.01; ***: p-value < 0.001.

Many of the genes which are lowly-expressed in GSE133383 (Figure 4A; LIM2, SPTSSB, LGALS9B, PRSS57, ADGRG3, and, to a lesser extent, DRAXIN), which lacks intestinal NK cells samples, are highly expressed in particular subsets of CRC-infiltrating NK cells (Figure 3A-B), possibly reflecting a role for these genes as CRC-specific NK cell markers induced by the tumour microenvironment. It is noteworthy that the peripheral blood CD56^bright^ subset – the most abundant NK cell subset in humans - do not cluster together, supporting the idea that this signature is more CRC-specific than previously reported NK cell signatures derived from blood-based analyses.

Next, as it is expected that genes which are considered robust markers of a given cell type should have coordinated expression patterns, the correlation between the individual genes in the signature was assessed. Correlation analysis demonstrated that the signature genes cluster into two major blocks where expression is highly cross-correlated, representing the genes selective for CD56^dim^ or CD56^bright^ NK cells (Figure 4B; orange and purple annotations at left). Subtype selectivity for each gene was defined based on its relative expression in each of these subsets (Figure 4C, Figure S6). That the two major blocks are anti-correlated may reflect the progressive loss of the CD56^bright^ markers as NK cells mature and reinforces the idea that our signature allows for pan-NK cell rather than subset-specific detection. Situated between the two major clusters are a subset of genes (including GNLY, TXK and PTGDR) which appear to have equivalent expression in CD56^bright^ and CD56^dim^ cells (Figure 4B; green annotations), likely corresponding to a “transitional” NK cell phenotype reported by several groups (52–54,57).

As a final visualisation measure, the expression of the NK cell signature in aggregate (ie. profiling expression of the signature as a whole, rather than the expression of each individual gene) was performed in an independent human dataset, GSE22886 (58). This dataset is composed of twelve different leukocyte subsets isolated from the PBMCs of healthy donors and was used to avoid the biases of testing our signature on the GSE60424 and GSE107011 data from which it was partially derived. In GSE22886, our NK cell signature was clearly enriched in the NK cell samples relative to the other cell types (Figure 5A). Due to the high sequence and transcriptomic homology between human and murine NK cells (59), we next interrogated the performance of the human NK cell signature in GSE15907 (Immgen; (60); a microarray dataset generated following the *ex vivo* isolation of multiple immune lineages from adult B6 mice (Figure 5B). Notably, in the murine context our NK cell signature clearly discriminates NK cells from CD8+ T-cells, γδ-T-cells and NKT cells, highlighting the flexibility of this novel NK cell signature to be used in *in vivo* studies to detect either endogenous murine or xeno-transplanted human NK cells.

**Figure 5.**
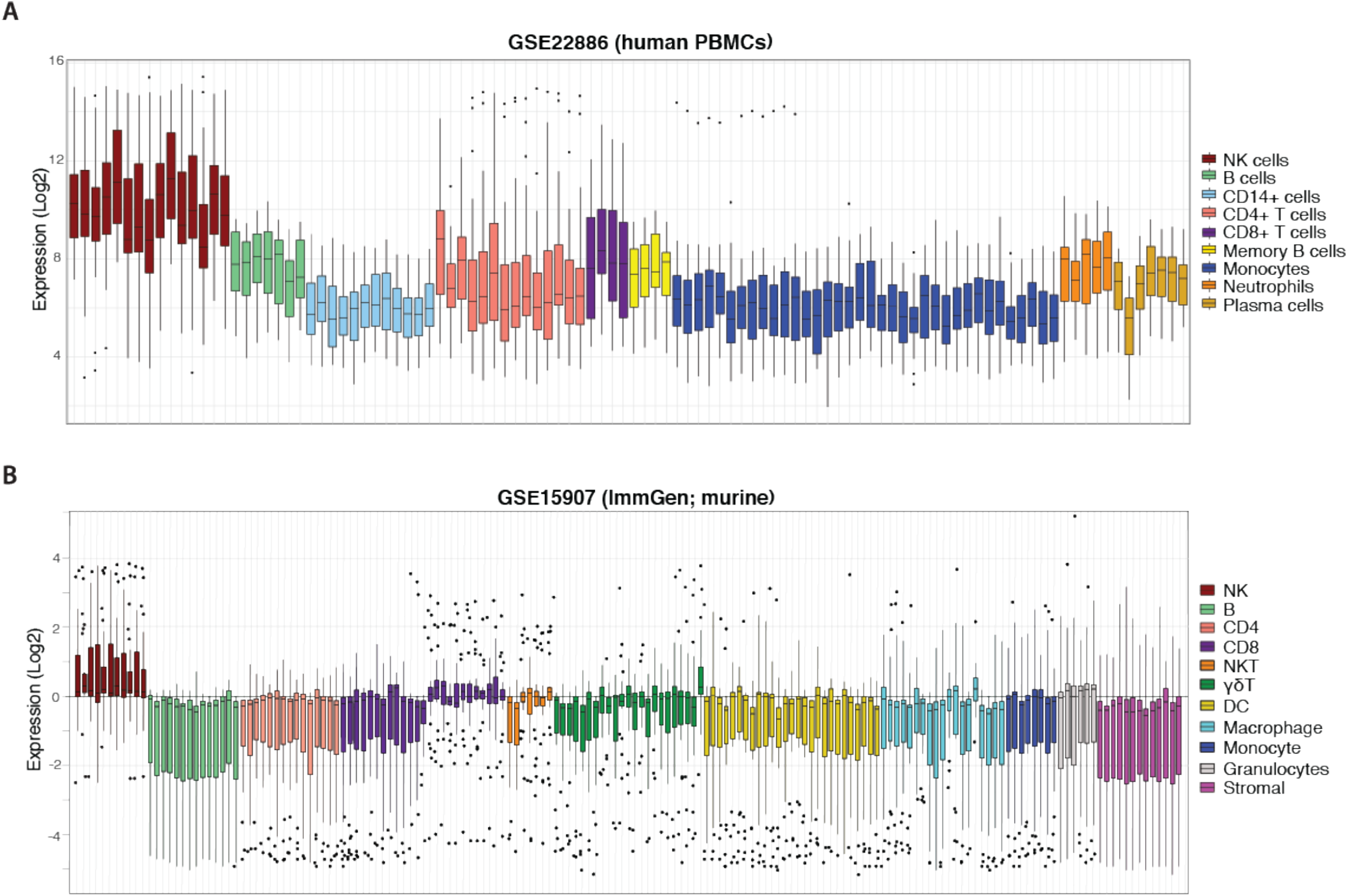
NK cell signature efficiently discriminates NK cells from other haematopoietic compartments. Boxplots of NK cell signature expression in sorted peripheral blood cells from **(A)** GSE22886 (Log2) independent human PBMC data and **(D)** GSE15907 (Immgen) murine data. Each boxplot represents one sample (coloured by cell type, at right) and individual points are single signature genes.

### Differential gene expression analysis confirms that cytotoxic and migratory programs are associated with high NK score

Having finalised the NK cell signature (Table S1, n_genes_ = 43), we next sought to determine which genes were concomitantly up- or downregulated in the samples whose transcriptomic profiles showed strong evidence of this signature. To identify these samples, we used a single-sample, rank-based gene-set scoring method termed *singscore* (42) to score samples against our NK signature. Here, a sample with a high NK score is interpreted as having high evidence of NK cell activity, whereas a sample with low NK score exhibits limited NK cell signature expression. Using samples sourced from two large, publicly available repositories of primary CRCs, the TCGA colorectal adenocarcinomas (TCGA-COAD; n_samples_ = 454), and GSE39582 (n_samples_ = 566, (62), we defined the NK-high and NK-low groupings based on the top 10% and bottom 10% of scored samples, respectively. We then performed differential expression analysis to compare the gene expression profiles (GEPs) of samples with high scores to those with low scores in each of the two data sets.

A strong positive correlation was observed between the logFC of genes in TCGA and GSE39582 datasets (Figure 6A-B; Spearman correlation ρ = 0.72, *p*-value < 2.2 x 10^-16^), suggesting similar transcriptional programs of NK inclusion/exclusion in both data sets.

**Figure 6.**
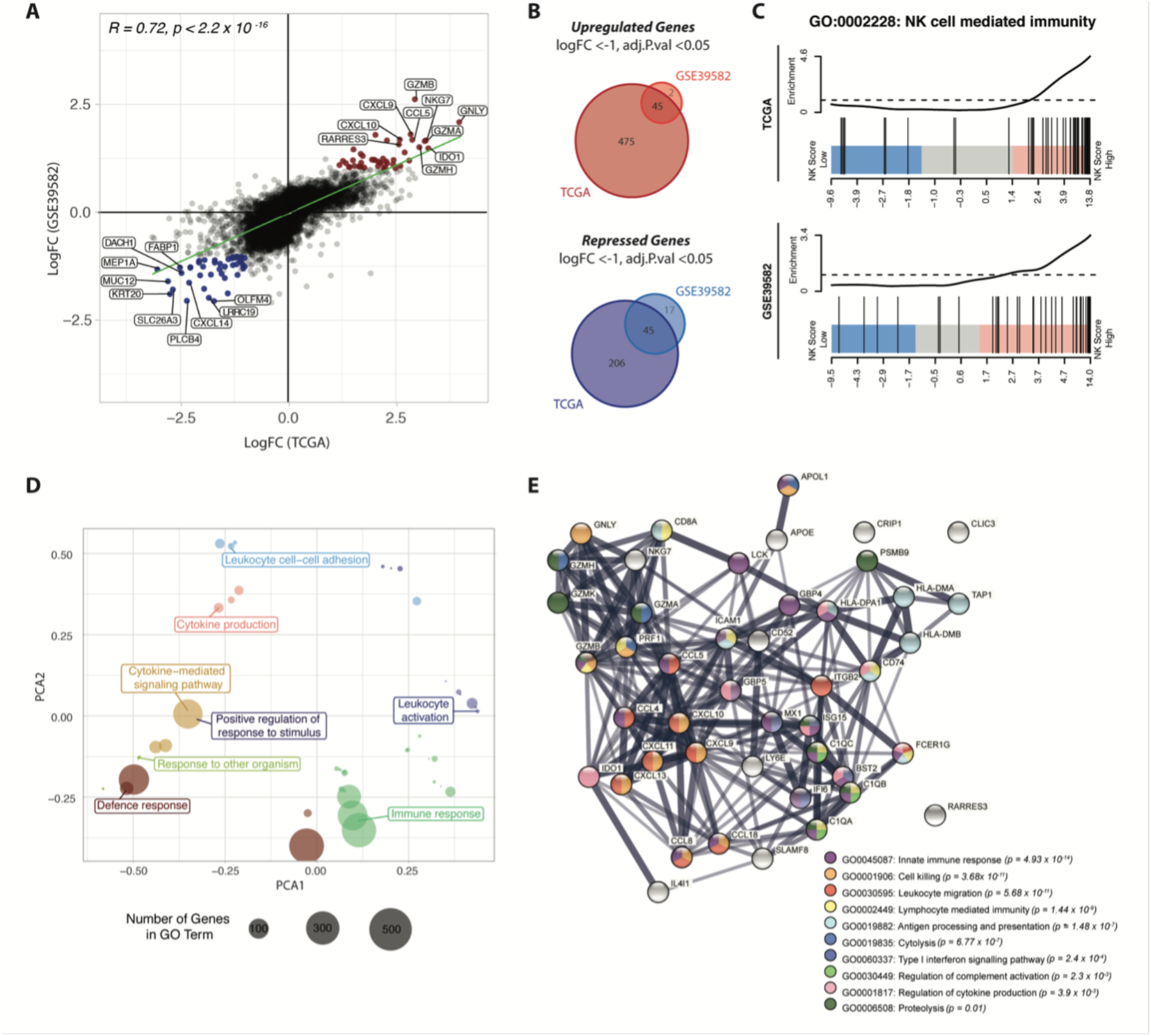
NK score is associated with high chemokine and cytolytic activity. **(A)** Scatterplot of the LogFCs of genes comparing NK-high and NK-low groups in TCGA and GSE39582 datasets; Genes that were differentially upregulated (red points) or repressed (blue points) in both datasets are highlighted. DEGs were defined as those with adjusted p-value < 0.05 and absolute LogFC > 1 or LogFC < −1. **(B)** Venn diagram of upregulated (*upper*) and down-regulated (*lower*) DEGs identified in (A). **(C)** Barcode plots showing enrichment of genes in the “NK cell mediated immunity” gene set (GO:0002228) in NK-high samples in both data sets **(D)** PCA plot of top 50 most significant GO terms in the NK-high group, distributed by semantic similarity of genes within each term **(E)** STRING network analysis of proteinprotein interactions between the 45 overlapping upregulated DEGs from (A), coloured by biological process. Thickness of the connecting line indicates the strength of evidence for the predicted interaction. *GO, gene ontology*

45 genes were differentially upregulated in NK-high samples from both datasets (Figure 6B, upper panel, Table S2). Of these, there was a strong enrichment of genes encoding cytotoxic effectors which are critical for NK cell killing, including the granule proteins NKG7 and granulysin (GNLY), as well as multiple members of the granzyme family including GZMB, GZMA and GZMH. Additionally, there was an overrepresentation of genes encoding ligands for chemokines implicated in NK cell trafficking, including CXCL9, CXCL10 and CCL5 (RANTES). Importantly, these ligands are expressed by tumour cells rather than the NK cells, suggesting that the high NK cell density in these tumours is at least partly driven by a tumour-intrinsic factor.

Conversely, 45 genes conserved between the two data sets were differentially repressed in the NK-high group (Table S3.3). The parallel downregulation of multiple genes which have been associated with increased metastasis and poor prognosis in CRC was observed, including Dachshund family transcription factor 1 (DACH1) (63) and the metalloprotease meprin-a (MEP1A) (64) and CXCL14 (46). Interestingly, recent evidence indicates that CXCL14 is essential for MHC-I upregulation (66) and as such, it stands to reason that CRC samples with high NK cell activity show decreased CXCL14 expression.

To identify biological processes associated with the NK-high phenotype, gene ontology (GO) enrichment analysis was conducted. Competitive gene set testing against the whole transcriptome of NK-high vs NK-low samples confirmed significant enrichment of the GO term “NK cell mediated immunity” in the NK-high group (Figure 6C). Moreover, the top 50 most significantly enriched GO terms in the NK-high group, when clustered by semantic similarity, converged on umbrella terms including “immune response”, “leukocyte activation”, “leukocyte cell-cell adhesion” and “cytokine production” (Figure 6D).

To visualise the functional synergy of the 45 overlapping upregulated DEGs (from Figure 6B) which define the NK-high group, the Search Tool for Retrieval of Interacting Genes/Proteins (STRING) platform was used to construct a protein-protein interaction (PPI) network (Figure 6E). Consistent with DE results, the PPI network was primarily centred around CC- and CXC-family chemokine ligands (CCL5, CCL4, CCL8, CXCL9, CXCL10, CXCL11, CXCL13, CXCL18) as well as cytolytic effectors (GNLY, GZMH, GZMK, GZMB, GZMK, NKG7, PRF1). A minor node related to antigen processing and presentation (HLA-DMA, HLA-DMB, HLA-DPA1, TAP1, CD74) may point towards a concomitant activation of effector T-cells in the NK-high group; however, chronic antigen stimulation is also known to induce lymphocyte exhaustion, resulting in reduced cytotoxic function (67).

Collectively, these data indicate that the NK-high group is defined by strong chemotactic signalling and high cytolytic activity. As these genes are signposts of an immune-active microenvironment, their upregulation in the NK-high group suggests that the NK cell signature selects for functionally active NK cells, rather than merely NK cell presence.

### High NK score is associated with improved survival outcomes and other clinical parameters

Given that NK cell load has been associated with better patient prognosis in CRC (21,23), we next performed Kaplan-Meier survival analysis to determine whether signature expression was associated with survival probability. Due to significant survival differences between patients with a primary or metastatic disease, we focussed only on those patients with Stage I-III CRC. For all survival analyses, patients were stratified by NK score where “NK-High” and “NK-Low” are defined as samples above and below the median NK score, respectively.

For the TCGA-COAD cohort, the NK-High group had significantly increased disease-free interval (DFI; defined as the period from date of diagnosis until a tumour progression event e.g. locoregional recurrence or distant metastasis) as compared with the NK-Low group (Figure 7A, log rank p-value = 0.0054, p-value from multivariate Cox regression model [accounting for age, stage and MMR status] = 0.02). Similarly, the GSE39582 cohort exhibited a trend towards significantly prolonged recurrence-free survival (RFS), defined as the period between surgical resection and a tumour progression event, at the univariate level (Figure 7B, log rank p-value = 0.053). There were no significant differences in overall survival (OS; Figure S7A, S7C) nor progression-free interval (PFI; Figure S37B) between the NK-High and NK-Low groups in either dataset.

**Figure 7.**
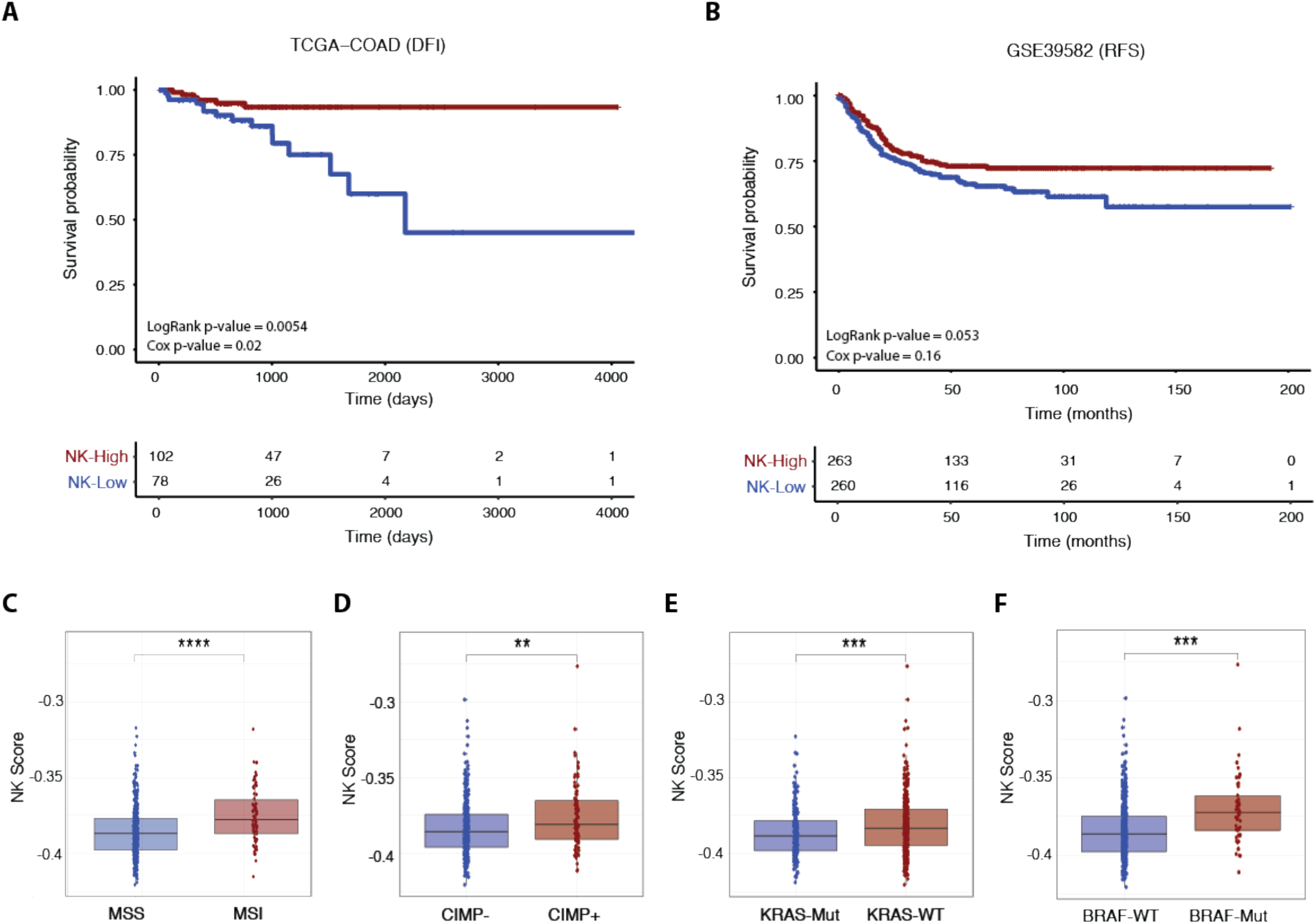
NK score is associated with survival outcome and other clinical parameters. Kaplan-Meier survival curves for patients stratified by NK score (where “NK-High” and “NK-Low” are defined as samples above and below the median NK score, respectively) for Stage IIII CRC patients in the **(A)** TCGA-COAD (DFI) and **(B)** GSE39582 (RFS) cohorts. Survival differences were tested using both log-rank and multivariate Cox proportional hazards models (adjusted for age, tumour stage and MMR status) with corresponding p-values indicated. **(C)** Boxplots of association of NK score with MSI **(D)** CIMP **(E)** KRAS and **(F)** BRAF status in GSE39582 patients (Student’s T-test; *: p-value < 0.05; **: p-value < 0.01; ***: p-value < 0.001). *DFI*, *disease-free interval; RFS, recurrence-free survival, MMR, mismatch repair; CIMP, CpG island methylator phenotype*.

We next interrogated the relationship between NK score and various clinical and molecular parameters such as patient history, molecular subtype, and driver mutation status. In both the GSE29582 (Figure S8A) and TCGA (Figure S9A) cohorts, NK scores were increased in early-stage disease (Stage I-II) as compared with late-stage disease (Stages III-IV). In the larger patient cohort, GSE29582, high NK score was significantly increased in those patients with MSI (Figure 7C) and CpG Island Methylator Phenotype (CIMP+; Figure 7D) disease. This was corroborated in the TCGA data (Figure S9A-B), where high NK score was also significantly enriched in those with MSI-associated clinical parameters such as tumour hypermutation (Figure S9D) and MLH1 silencing (Figure S9E). With respect to driver mutations, NK score was significantly enriched in those patients with KRAS wildtype (Figure 7E) or BRAF mutant (Figure 7F) genotypes, although there was no association with TP53 status (Figure S8D).). There was no association between NK score and tumour characteristics such as tumour location (Figure S8B; distal versus proximal colon) and histological subtype (Figure S9F; mucinous vs non-mucinous). Similarly, no association was found between NK score and factors related to clinical history including adjuvant chemotherapy status (Figure S8C), prior CRC diagnosis (Figure S9G) or evidence of synchronous disease (Figure S9H). In sum, these results support the clinical utility of using this newly derived NK cell signature in the context of CRC.

## Discussion

Bioinformatic approaches which allow for the deconvolution of immune cell subsets from bulk sequencing data have revolutionised our ability to assess the immune landscape of individual tumours. Transcriptomic signatures which predict intra-tumoral NK cell infiltration have been shown to indicate improved patient survival in various cancers (40,41). However, despite accumulating evidence that NK cell load is prognostic in CRC (21,23), there are currently no extensively curated NK cell signatures explicitly designed for use in the context of CRC.

Here, through extensive computational curation of established NK cell-related genes with putative markers discovered through DE analysis of various bulk RNAseq and scRNAseq datasets, we define a comprehensive NK cell signature specific for CRC samples. Survival modelling in two large cohorts of primary CRC patients indicated that NK signature expression is associated with prolonged progression- and recurrence-free intervals, consistent with previous reports on the beneficial prognostic impact of NK cell infiltration in other solid cancers (68). That these metrics are associated with tumour progression (rather than overall survival) may relate to the fact that NK cells are believed to play a greater role in the suppression of metastases rather than the prevention of primary tumours (16,18,69–71).

Although the NK cell signature presented herein is not the first of its kind, we believe that it is the most specific and the most extensively curated NK cell signature for the precise identification of the wide range of NK cell subsets in CRC samples due to its derivation in the context of this tumour type. Practical limitations have meant that most NK cell gene signatures, including that employed in the widely used CIBERSORT immune cell deconvolution tool, were curated using NK cells isolated from the peripheral blood. As NK cells exhibit tissue-specific phenotypes (37,52,72), and the tumour microenvironment is known to rewire NK cell transcriptional programs (72–76), it is difficult to assess how accurately these pre-existing NK cell signatures perform when extrapolated for use in the context of cancer. Moreover, striking differences in the transcriptomic profiles of circulating NK cells versus tumour-infiltrating NK cells from the same patients have also been reported (76), emphasising the importance of using scRNAseq data from tumour-infiltrating NK cells to faithfully derive a reference transcriptomic profile for CRC-associated NK cells.

Another issue facing pre-existing NK cell signatures has been a potential lack of specificity. For example, although a four-gene signature (composed of NCR1, PRF1, CX3CL1 and CX3CR1) was shown to accurately distinguish NK-high vs NK-low subgroups in clear cell renal cell carcinoma (41), these genes are ubiquitously expressed by many immune cell types. More broadly, the transcriptional programs of NK cells are highly overlapping with those of multiple T-cell subsets, particularly γδ-T-cells (77). Numerous studies have demonstrated that several archetypical NK cell markers including NKG2A (78), NCRs (79,80) and various KIRs (81) are expressed by both traditional αβ- and unconventional γδ-T-cells. Our strategy of prioritising genes based on their NK-cell specificity rather than only those with the highest expression may therefore increase precision when teasing apart the NK cell contribution from complex mixtures of other immune cells.

In both the TCGA and GSE39582 cohorts investigated, NK scores were higher in patients with early-stage disease, aligning with results of a large, pan-cancer meta-analysis reporting that NK cell infiltration is lower in advanced-stage tumours (68). Similarly, high NK score was associated with several clinically useful molecular parameters such as MSI disease, CIMP positive status and BRAF mutation which have each been previously linked with high TIL and NK cell infiltration (82,83) and are also defining characteristics of the highly immunogenic CMS1 molecular subtype of CRC (84). Moreover, DE analysis between the NK high- and low-scoring patients identified transcriptional and biological processes associated with high cytotoxic and chemokine activity in samples with high NK scores. As the signature efficiently discriminates murine NK cells from other immune cell types, this signature may also prove useful in prospective *in vivo* studies.

It remains unclear how optimally transcriptional signatures validated in a particular cancer will perform in alternate tumour types. Thus, future work may focus on cross-validating NK cell signatures in contexts outside their tumour-of-derivation, and/or on defining a pan-cancer NK cell signature which is less sensitive to subtype-specific influences. Prospective studies may also aim to dissect the transcriptional profile of intra-metastatic NK cells and determine whether, or in what ways, this diverges from that of primary CRC-infiltrating NK cells.

