## Supplementary figures and images for "A new highly-specific Natural Killer cell-specific gene signature predicting recurrence in colorectal cancer patients"

### Supplemental Figures

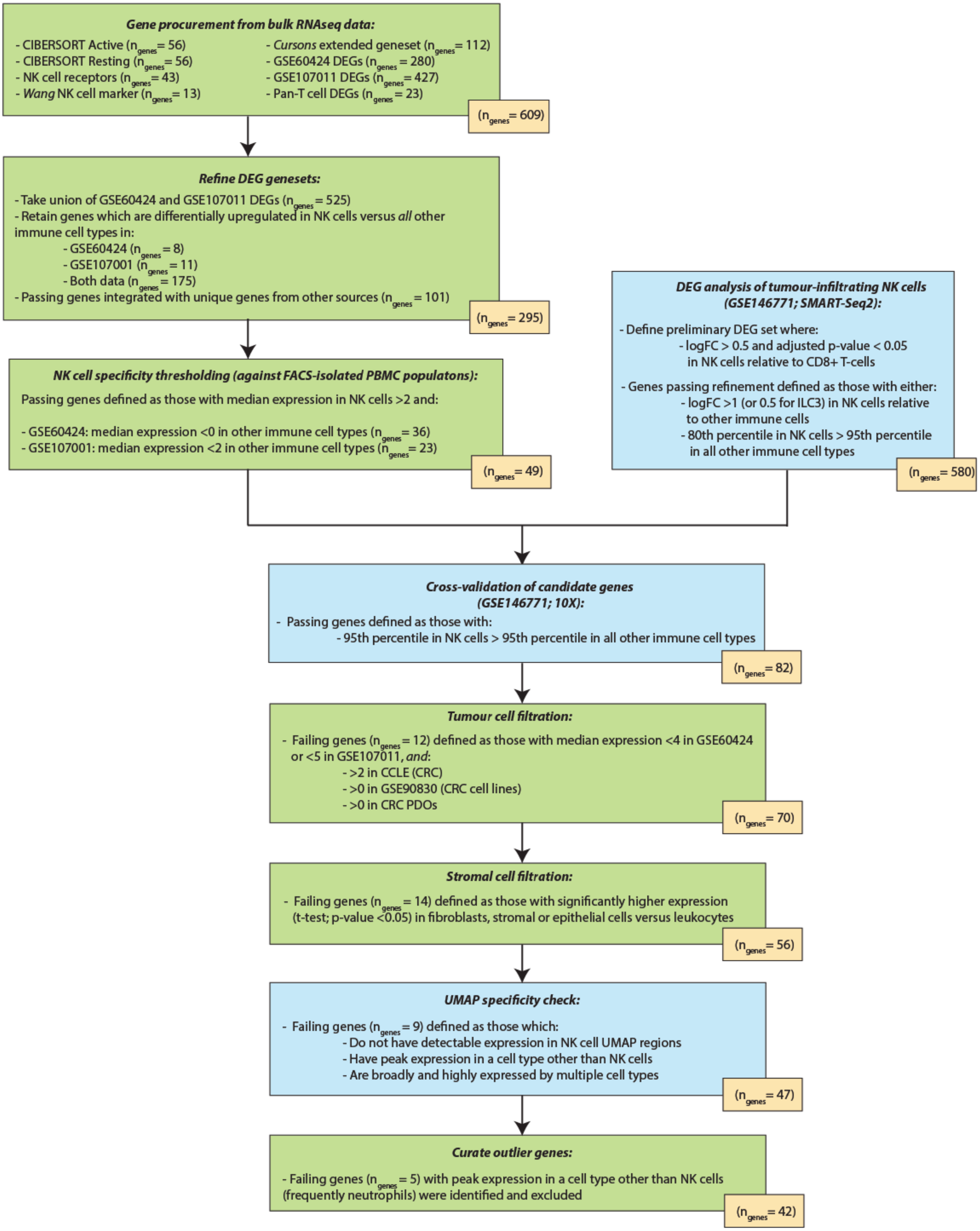

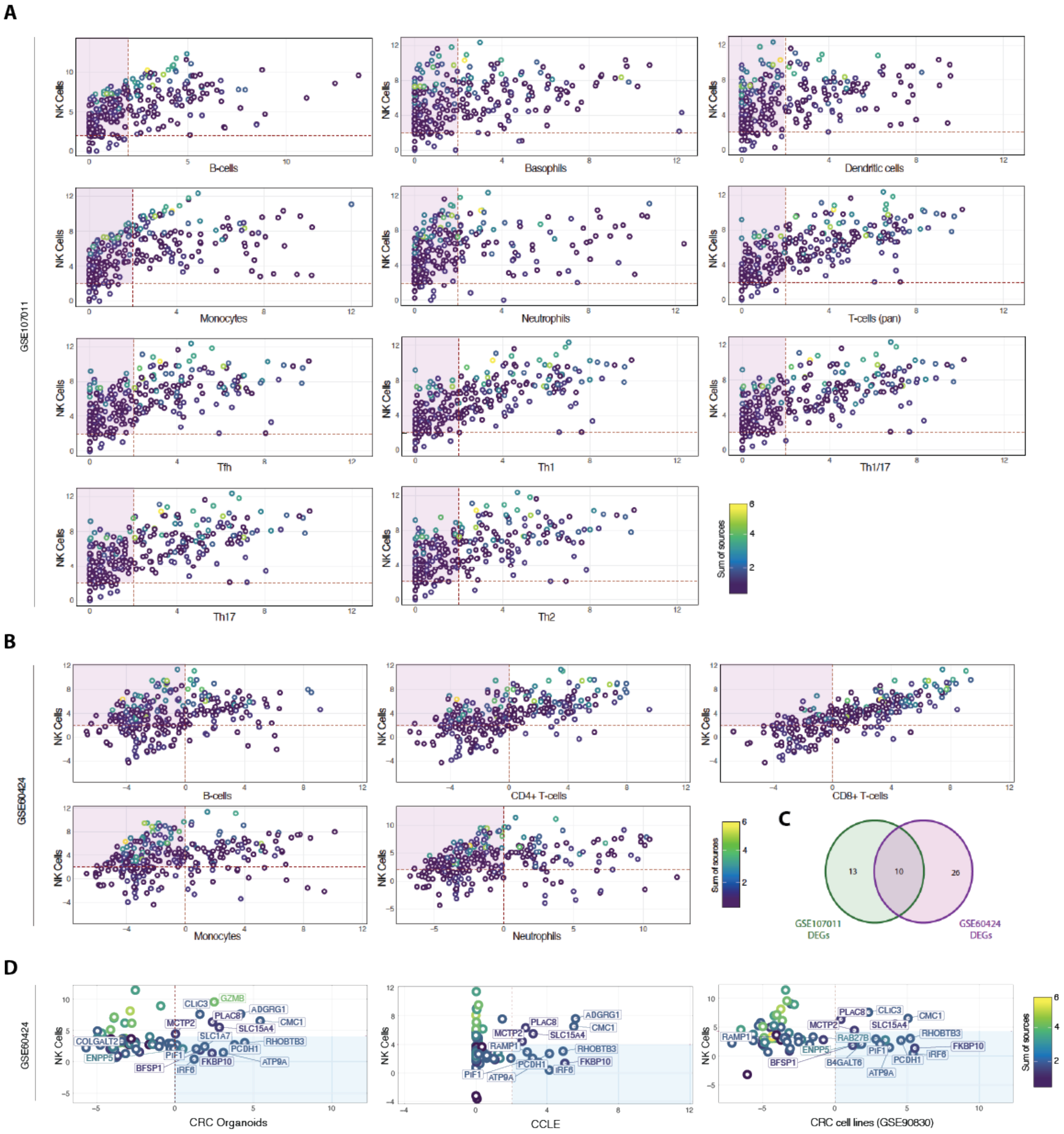

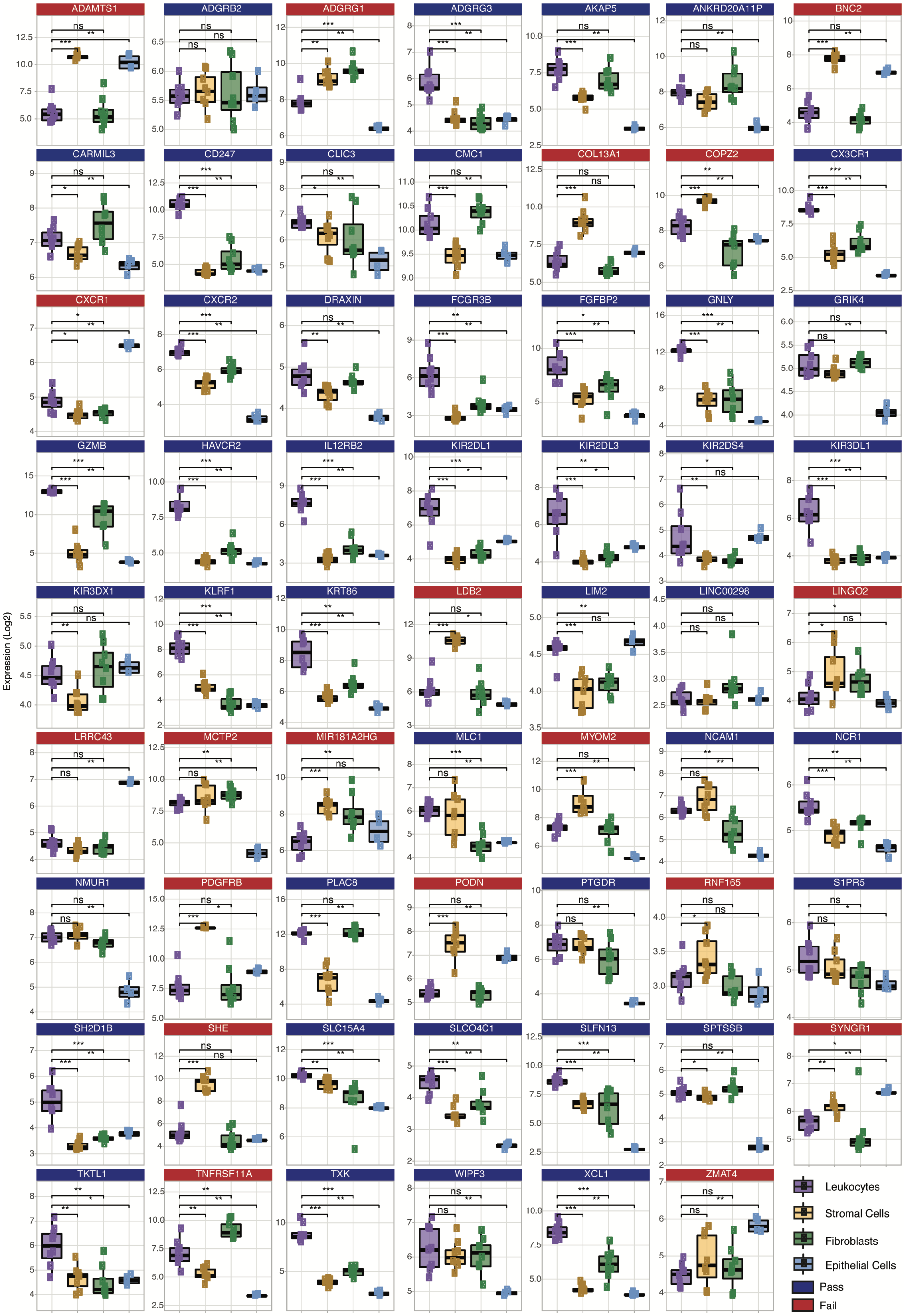

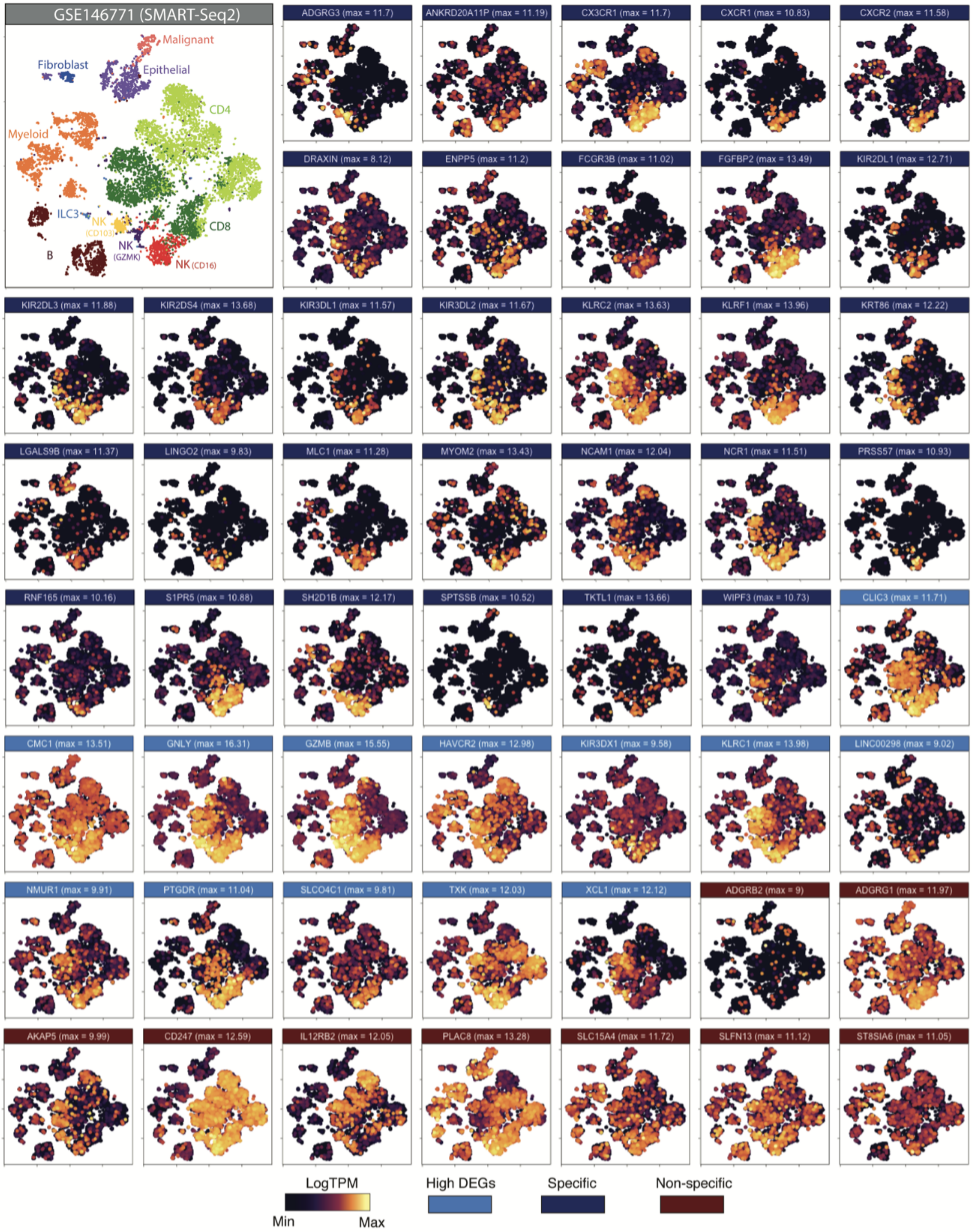

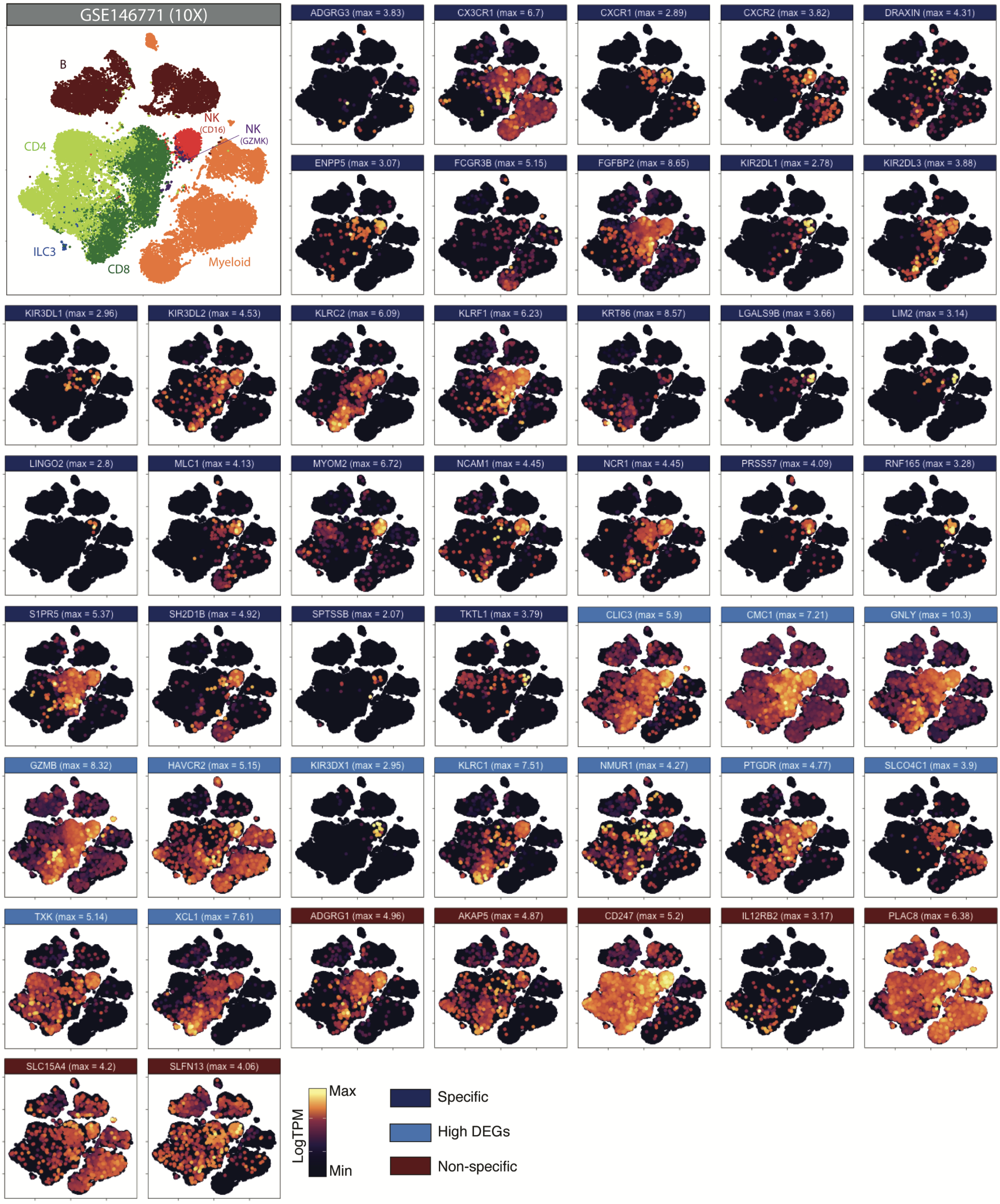

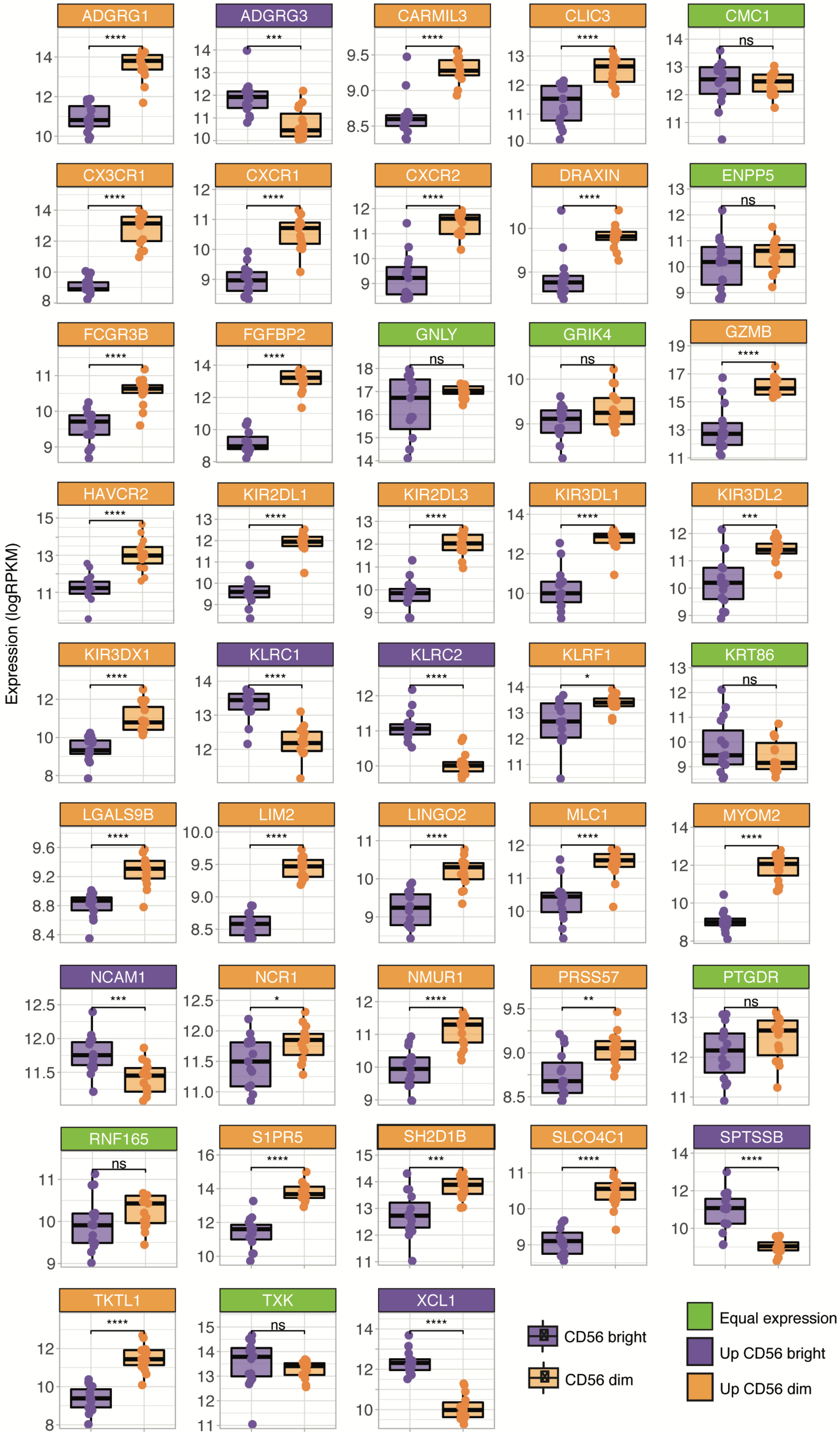

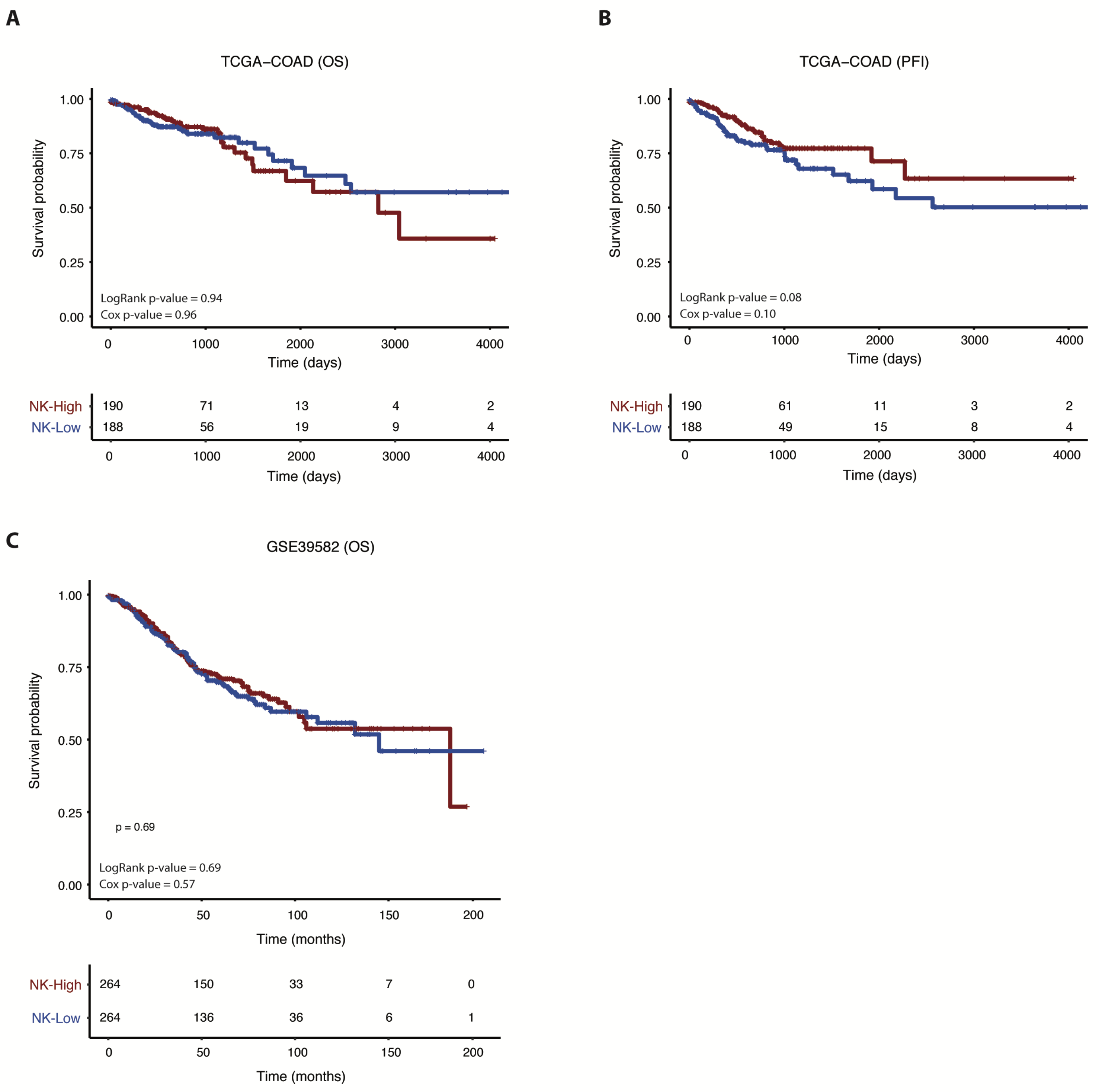

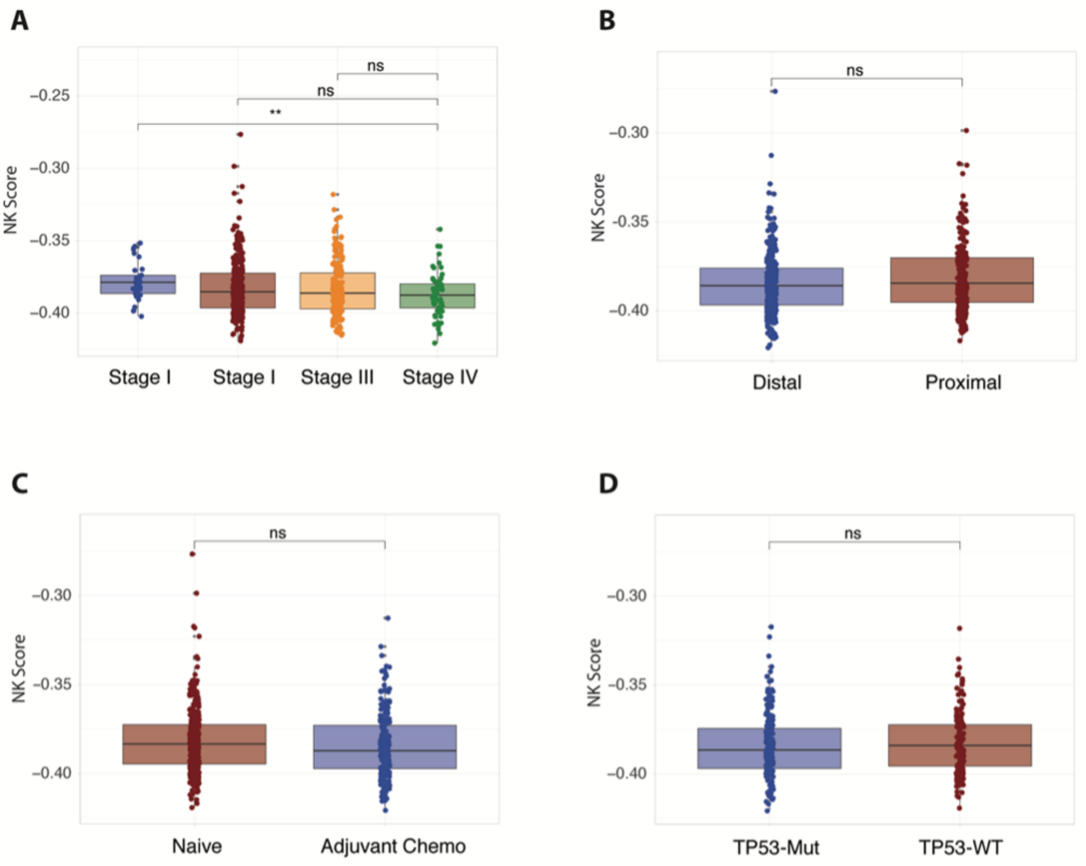

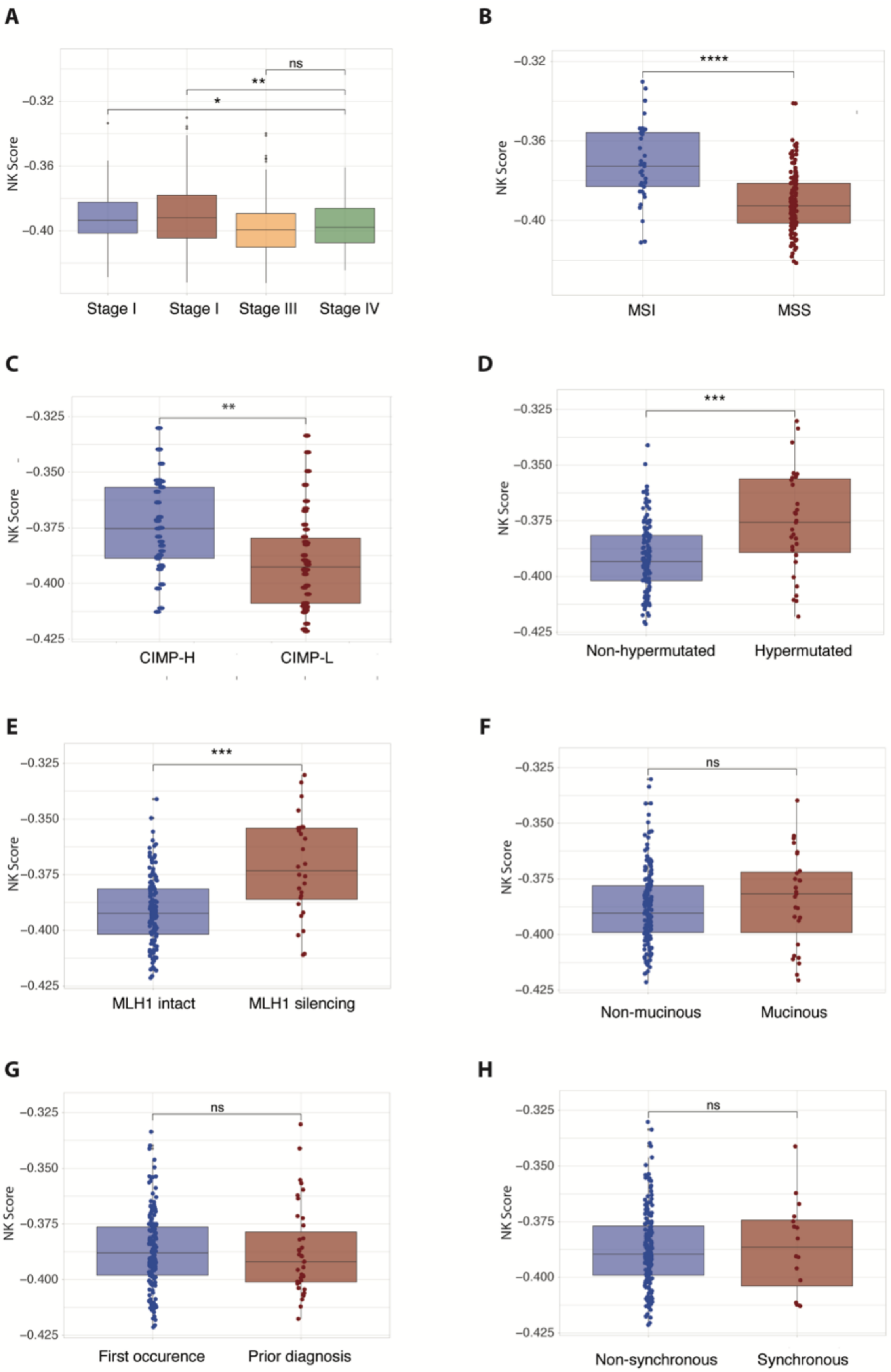
