## Supplemental Figure Legends for "A new highly-specific Natural Killer cell-specific gene signature predicting recurrence in colorectal cancer patients"

**Figure S1 | Overview of NK cell signature curation pipeline.** A preliminary gene set ($n_{genes}$ = 609) compiled from eight partially overlapping sources was sequentially filtered for NK cell specificity against immune, tumour and stromal cells in multiple bulk RNAseq (green boxes) and scRNAseq (blue boxes) datasets.

**Figure S2 |Additional NK cell specificity filtration against immune and CRC-supportive cell types.** Biplots depicting median expression for each candidate gene (rings; coloured by sum of sources) in NK cells from **(A)** GSE60424 (logRPKM) and **(B)** GSE107011 (logTPM) versus other immune cell types. The intersect of passing genes for each pairwise comparison (purple boxes) were retained as candidate genes. **(C)** Venn diagram of candidate genes retained from (A) and (B). **(D)** Biplots depicting median expression for each candidate gene (rings; coloured by sum of sources) in NK cells from GSE60424) versus CRC organoids (left column), CCLE CRC cell lines (centre column) and GSE90830 (CRC cell lines; right column). The union of failing genes for each pairwise comparison (blue boxes) were flagged for removal from the candidate geneset.

**Figure S3 |Candidate gene filtration against CRC-associated stromal cells.** Boxplots of candidate gene expression in the leukocyte (CD45+; purple), stromal cell (CD31+; gold), fibroblast (FAP+; green) and epithelial cell (EpCAM+; light blue) fractions isolated from the tumours of 6 patients with CRC (GSE39396). Failing genes (red headers) were defined as those with significantly higher expression relative to leukocytes in a non-leukocyte subset (Student's T-test; *: p-value < 0.05; **: p-value < 0.01; ***: p-value < 0.001) whereas passing genes (dark blue headers) were those significantly enriched in leukocytes relative to the other cell types or those expressed uniformly across all four cell types.

**Figure S4 | Candidate gene expression in CRC-infiltrating cellular subsets.** UMAP plots of dissociated CRC samples (GSE146771; SMART-Seq2) coloured by cell type (top-left) and candidate gene expression. Candidate genes were classified as either NK cell specific (dark blue headers), "High DEGs" (light blue headers) or non-specific to NK cells (red headers; subsequently removed from candidate gene set). Maximum expression is indicated in parentheses.

**Figure S5 | Candidate gene expression in CRC-infiltrating immune cells.** UMAP plots of dissociated CRC samples (GSE146771; 10X) coloured by cell type (top-left) and candidate gene expression. Candidate genes were classified as either NK cell specific (dark blue headers), "High DEGs" (light blue headers) or non-specific to NK cells (red headers; subsequently removed from candidate gene set). Maximum expression is indicated in parentheses.

**Figure S6 | Comparison of refined gene set in $CD56^{bright}$ vs $CD56^{dim}$ NK cells.** Boxplots of refined gene set in NK cells isolated from the blood, spleen, bone marrow, lung and lymph nodes of four healthy donors (GSE133383). Genes are annotated (headers) according to whether they show significant upregulation in the $CD56^{dim}$ (orange), $CD56^{bright}$ (purple) or neither (green) (Student's T-test; *: p-value < 0.05; **: p-value < 0.01; ***: p-value < 0.001).

**Figure S7 | Additional survival analyses in TCGA-COAD and GSE39582 data.** Kaplan-Meier survival curves for patients stratified by NK score (where "NK-High" and "NK-Low" are defined as samples above and below the median NK score, respectively) for Stage I-III CRC patients in the **(A)** TCGA-COAD (OS), **(B)** TCGA-COAD (PFI) and **(C)** GSE39582 (OS) cohorts. Survival differences were tested using both log-rank and multivariate Cox proportional hazards models (adjusted for age, tumour stage and MMR status). *OS, overall survival; PFI, progression-free interval, MMR, mismatch repair.*

**Figure S8 | Association of NK score with clinical parameters in GSE39582 data.** Boxplots showing association of NK score with **(A)** tumour stage **(B)** TP53 status **(C)** adjuvant chemotherapy status and **(D)** tumour location in GSE39582 patients (Student's T-test; *: p-value < 0.05; **: p-value < 0.01; ***: p-value < 0.001).

**Figure S9 | Association of NK score with clinical parameters in TCGA-COAD data.** Boxplots showing association of NK score with **(A)** tumour stage **(B)** MMR status **(C)** CIMP status **(D)** hypermutation status **(E)** MLH1 silencing status **(F)** histological subtype **(G)** clinical history of CRC and **(H)** presence of synchronous disease in TCGA-COAD patients (Student's T-test; *: p-value < 0.05; **: p-value < 0.01; ***: p-value < 0.001).
